# Two-photon volumetric study of cleared, invasive ductal carcinoma breast tissue samples and associated axillary lymph nodes

**DOI:** 10.1101/2025.11.07.687157

**Authors:** Uma Pisarović, Loes F.S. Kooreman, Thiemo J.A. van Nijnatten, Albert Bitorina, Sven Hildebrand, Anna Schueth

## Abstract

**Background:** Breast cancer remains the most frequently diagnosed cancer among women worldwide, with invasive ductal carcinoma (IDC) representing the most common subtype. Despite its high prevalence, current diagnostic workflows rely on 5 µm-thin haematoxylin and eosin-stained (H&E) sections, inherently limiting spatial insight into tumor architecture and extracellular matrix (ECM) organization. As the interest in studying intratumoral heterogeneity increases, three-dimensional (3D) imaging is becoming an increasingly valuable tool.

**Methods:** This study applied a tissue clearing and imaging pipeline to large formalin-fixed paraffin-embedded (FFPE) breast and lymph node tissue using IDC samples from Maastricht University Medical Centre+. Deparaffinized samples were processed using a modified MASH protocol. Tissues were labelled with Neutral Red, Eosin Y, Methyl Green, and DAPI. Tissue shrinkage was analysed across all processed samples. Two-photon (2P) microscopy was used to image malignant and non-malignant breast tissue, and matched axillary lymph nodes from a patient with grade I IDC to depths of up to 1000 µm via DAPI and second harmonic generation (SHG) channels. Image analysis included assessments of dye penetration, nuclear and collagen content, and fiber orientation using FIJI software.

**Results:** Clearing and staining preserved tissue structure and achieved high transparency across millimetre-scale volumes. 2D surface shrinkage averaged 6.7% (p < 0.001). DAPI signal penetration was consistent with SHG signal profiles up to approximately 600 µm of tissue depth. Structures such as terminal ductal lobular units, adipocytes, vasculature, and lymphoid follicles were clearly visualized in 3D. Quantitative and qualitative analysis in grade I IDC tissue revealed regional differences in cell size and shape, collagen content, and fiber coherency, indicating localized early-stage ECM remodelling.

**Conclusion:** This is the first study to apply 2P microscopy with MASH clearing to large FFPE IDC breast and lymph node samples. The protocol enables reproducible high-resolution volumetric imaging and lays the groundwork for future research applications, leading to potential diagnostic applications.

## Introduction

In 2022, breast cancer was the most commonly diagnosed cancer among women, accounting for 26.7% of all cases. It also represented the highest number of new diagnoses globally, with 2,296,840 cases (23.8%), and was responsible for 666,103 cancer-related deaths (15.4%) [1]. Pathologically, breast cancer is classified based on anatomical origin, invasiveness, and molecular subtype. Most tumors arise from the terminal duct lobular units (TDLUs) and are categorized as either ductal or lobular carcinomas. They are further divided into non-invasive (in situ) or invasive types depending on their capacity to breach the basement membrane. The most commonly diagnosed forms include ductal carcinoma in situ (DCIS), lobular carcinoma in situ (LCIS), invasive lobular carcinoma (ILC), and invasive ductal carcinoma (IDC), also referred to as no special type or not otherwise specified, with IDC accounting for approximately 80% of all invasive cases [2–4].

Histological grading of IDC, typically performed according to the Nottingham modification of the Bloom–Richardson system, evaluates three morphological features: tubule formation, nuclear pleomorphism, and mitotic activity, each scored on a scale from 1 to 3. The combined score classifies tumors as Grade I (well-differentiated), Grade II (moderately differentiated), or Grade III (poorly differentiated), correlating with increasing tumor aggressiveness and poorer clinical outcome [5]. This grading is performed on hematoxylin and eosin (H&E)-stained formalin-fixed, paraffin-embedded (FFPE) sections, which remain the clinical gold standard. However, these 5 µm thin sections provide only narrow, two-dimensional snapshots of the complex tissue architecture in volumetric tumors. Such thin sections cannot capture intratumoral heterogeneity, a major determinant of treatment response, disease progression, and clinical outcome [6,7]. Limited sampling of a few isolated planes may overlook spatial variations, and some pathological features may go undetected. Scientific findings based on large-volume tissue information are therefore more broadly applicable.

Addressing these limitations requires imaging approaches capable of high-resolution, three-dimensional (3D) visualization of tissue microarchitecture. Two-photon (2P) microscopy provides a powerful solution, offering deep tissue penetration and high spatial resolution without physical slicing, enabling exploration of intact tumor volumes across hundreds of microns [8,9].

Several studies have applied multiphoton microscopy (MPM) in two-dimensional settings [10–14]. By combining two-photon excited fluorescence (TPEF) and second harmonic generation (SHG), MPM enables high-resolution imaging without exogenous dyes or extensive sample preparation. In breast tissue, endogenous molecules such as nicotinamide adenine dinucleotide (phosphate) (NAD(P)H) and flavin adenine dinucleotide (FAD) produce TPEF signals reflective of metabolic activity, while collagen fibers generate strong SHG signals [15]. This allows simultaneous imaging of cellular metabolism and extracellular matrix (ECM) organization.

Burke et al. used SHG imaging on 5 µm FFPE sections to monitor collagen structural changes across healthy tissue, DCIS, LCIS, IDC, and ILC, and found significant alterations between healthy and malignant tissue [10]. Wu et al. performed MPM on fresh 6 µm paired cancerous and healthy slices from 20 IDC patients, revealing increased nuclear density and collagen disorganization in tumors [11]. Natal et al. applied SHG to 60 FFPE luminal B breast cancer samples and showed that specific collagen features were predictive of prognosis [12]. Han et al. compared MPM imaging to immunohistochemistry (IHC) and H&E in DCIS and IDC samples, demonstrating higher specificity with fewer preparatory steps than IHC [13]. Dinkle et al. used SHG microscopy on 5 µm FFPE sections from triple-negative breast cancer to characterize collagen alterations linked to tumor aggressiveness [14].

Despite these promising results, all of these studies were limited to thin 2D sections, making it difficult to assess the spatial complexity of tumor architecture and ECM interactions fully. 3D applications of MPM have been explored, but largely in breast cancer spheroids [16] or mouse mammary tissue [17]. A notable knowledge gap remains regarding whole FFPE human breast tissue blocks and 3D IDC architecture. Moreover, volumetric investigation of associated lymph node tissue is largely absent in current research.

In the present study, we introduce an adaptation of a clearing and imaging pipeline tailored for large (millimeter-scale) FFPE human breast and lymph node tissue specimens, including matched malignant, adjacent non-malignant, and lymph node tissues from IDC patients. This approach is based on the previously described MASH (iDISCO) clearing protocol, combined with 2P microscopy to evaluate imaging depth, structural preservation, and quantitative differences across malignant, healthy, and lymph node tissues. Nuclear and collagen content, fiber orientation, and prominent tissue structures were assessed. To our knowledge, this is the first application of MASH in large-scale human FFPE breast and lymph node samples using two-photon imaging, demonstrating the feasibility and added value of volumetric imaging for future research and potential diagnostic applications [18,19].

## Materials and Methods

### 2.1 Human FFPE breast and lymph node tissue

FFPE human tissue specimens were obtained from the Department of Pathology at Maastricht University Medical Centre+ (MUMC+) following surgical resection and standard histopathological processing. Tissues were collected from three patients diagnosed with IDC, comprising malignant breast tissue, adjacent non-malignant breast tissue, and matched axillary lymph nodes from each individual. All specimens were fully anonymized and handled in accordance with the propositions of the Medical Ethics Review Committee of MUMC+ in the Netherlands.

### 2.2 Deparaffinization of human FFPE breast and lymph node samples

Deparaffinization of FFPE samples was performed by incubating each specimen in 50 mL of *paraffinum liquidum* for two days in a water bath preheated to 60°C, followed by sequential incubations in 50 mL of xylene at room temperature (RT) on a shaker for 1 hour and overnight, respectively. Following deparaffinization, the samples were rehydrated in descending concentrations of 50mL of ethanol in distilled water with 1-hour incubations at RT on a shaker in the following sequence: 100% ethanol (twice), 70%, and finally, 50%. To preserve the samples overnight, they were each incubated in 50mL of 0.1 M phosphate-buffered saline (PBS) at RT on a shaker.

### 2.3 Adapted protocol for clearing and labelling of deparaffinized human breast and lymph node samples

The clearing and labeling protocol was adapted from the MASH (iDISCO+) method previously developed and described by Hildebrand et al. [18]. Their novel MASH refractive index matching solutions (RIMS), adjusted to a refractive index (RI) of 1.56, were substituted with ECi, which provides comparable tissue transparency [20]. Additionally, dye combinations and working solutions have been adjusted. Similar as in the work performed by Hildebrand et al., the protocol consists of the following steps: (1) sample pretreatment with methanol and bleaching, (2) labeling with selected dyes as explained below, and (3) tissue clearing and refractive index matching. For pretreatment, bleaching and labeling the samples were kept and processed in 10mL six-well plates

#### 2.3.1 Sample pretreatment with methanol and bleaching

For the pretreatment process, samples were each dehydrated in 8mL of ascending concentrations of methanol (MeOH) diluted in distilled water. This involved sequential 1-hour incubations at RT on a shaker in 20%, 40%, 60%, and 80% MeOH, followed by two 1-hour incubations in 100% MeOH, with the second carried out at 5 °C in a cold room. Bleaching was performed overnight at 5 °C on a shaker using 8 mL per sample of freshly prepared 5% hydrogen peroxide solution, made by mixing one part 30% H₂O₂ with five parts MeOH. After re-equilibration to RT, samples were rehydrated through descending MeOH concentrations (60%, 40%, and 20%), each for 1 hour. This was followed by permeabilization in 8 mL per sample of 0.1 M PBS/0.2% (v/v) Triton X-100 at pH 7.4, with incubations for 1 hour and then overnight. All rehydration and permeabilization steps were conducted at RT on a shaker.

#### 2.3.2 Labelling with MASH dyes, Eosin Y, and counterstaining with DAPI

Before labeling, samples were incubated for 1 hour at RT, on a shaker in 8 mL per sample of freshly filtered aqueous solution of 50% (w/v) potassium disulfite. This was followed by five quick rinses in distilled water and a subsequent 1-hour incubation in 8 mL distilled water at RT on a shaker. The following dyes were used to label cytoarchitectural features, each prepared in phosphate-citrate buffer (McIlvain buffer) at pH 4.0: 0.001% neutral red (NR), 0.00025% methyl green (MG), and 0.00025% Eosin Y. For nuclear counterstaining, 0.001% DAPI in McIlvain buffer (pH 4.0) was used. Labeling was performed in 8 mL per sample of dye, over four days at RT on a shaker, following the scheme outlined in Table 1. Samples were flipped halfway through the incubation period to ensure uniform staining.

**Table 1:**
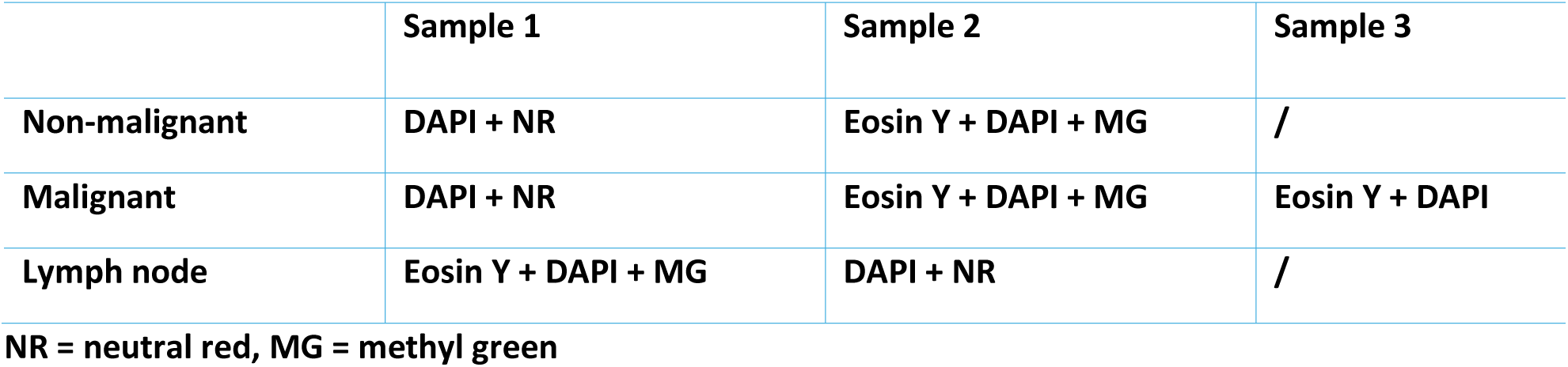
Dye combinations used for labeling of different tissue samples from patients with IDC.

|  | Sample 1 | Sample 2 | Sample 3 |
| --- | --- | --- | --- |
| Non-malignant | DAPI + NR | Eosin Y + DAPI + MG | / |
| Malignant | DAPI + NR | Eosin Y + DAPI + MG | Eosin Y + DAPI |
| Lymph node | Eosin Y + DAPI + MG | DAPI + NR | / |

NR = neutral red, MG = methyl green

#### 2.3.3 Clearing and refractive index matching with ECi

Following labeling, samples were washed twice for 1 hour in 8 mL per sample of McIlvain buffer (pH 4), then dehydrated through sequential 1-hour incubations in ascending concentrations of MeOH in distilled water (20%, 40%, 60%, 80%, and twice in 100%). Samples were subsequently transferred into 50 mL high-density polyethylene (HDPE) incubation tubes and incubated overnight in a 33% MeOH / 66% dichloromethane (DCM) solution for delipidation. Residual methanol was removed by two 1-hour incubations in 100% DCM. At each step, the 50 mL tubes were completely filled with the respective solution. Finally, samples were immersed in ECi as a RIMS. All procedures were conducted at RT on a shaker.

### 2.4 Two-photon imaging and visualization using Leica Application Suite Advanced Fluorescence (LAS AF) software

2P imaging was conducted on patient 1 (IDC grade I) malignant breast tissue, adjacent non-malignant breast tissue, and matched lymph node tissue. For this paper, a total of three malignant breast tissue volume stacks, two non-malignant breast tissue volume stacks, one lymph node volume stack, and one single-plane lymph node image were included in the analysis. Each volume stack was acquired through two channels, one capturing the DAPI signal and the other capturing second harmonic SHG signal. Before imaging, stained and cleared samples were placed onto a glass petri dish, and overlaid with a water droplet and a cover slip. A 2P laser scanning microscope (Leica TCS SP5 MP, Leica Mikrosysteme Vertrieb GmbH, Wetzlar, Germany), with a HCX APO L 20x/1.00 W water immersion objective was used. The excitation source was a laser with a wavelength set to 800 nm. To avoid photobleaching and tissue damage, laser power was kept at 11%. Images and image stacks were created and visualized in Leica Application Suite Advanced Fluorescence (LAS AF) software. Imaging parameters, including step size, zoom factor, scan speed, image resolution, and laser power, varied slightly between stacks and are detailed in Supplementary Table 1.

### 2.5 Data analysis using FIJI (ImageJ) and Microsoft Excel

Image analysis was performed using FIJI (ImageJ), while statistical analysis and graphical representation were carried out in Microsoft Excel 2019. Images were imported into FIJI via the Bio-Formats Importer. Individual channels were split, and each relevant grayscale image was processed separately according to the specific requirements of the analysis.

#### 2.5.1 Tissue Shrinkage Analysis

2D tissue shrinkage was assessed by comparing the surface area of samples before and after processing. The analysis was performed on all successfully processed samples, including malignant breast tissue, adjacent non-malignant breast tissue, and matched axillary lymph nodes from patient 1 and patient 2, as well as malignant breast tissue from patient 3. Progress photographs were taken during tissue processing with the samples placed on millimeter paper to provide a scale reference. These images were imported into FIJI, where the tissue was manually outlined as a region of interest (ROI), and the area fraction of the tissue relative to the total image was measured. To improve measurement reliability, each tissue sample was outlined and measured three times, and the mean area fraction was calculated. The absolute tissue area was then determined by multiplying the mean area fraction by the total image area, calculated based on the known dimensions of the mm paper. In cases where tissue was partially damaged or fragmented during processing, the initial area was measured without the separated portions to ensure consistency. A one-tailed paired-sample t-test with a significance level of α = 0.05 was used to evaluate whether the decrease in tissue area after processing was statistically significant. All area ROIs used for this analysis are shown in Supplementary Figures 1–3). Tissue depths were approximated visually using millimeter paper, however due to the imprecision of the method, and varying tissue thicknesses across the tissue area, the 3D tissue shrinkage analysis was not performed.

#### 2.5.2 DAPI Dye Penetration analysis

Signal intensity profiles of the DAPI (nuclear) channel and the SHG (collagen) channel were generated in FIJI for volume stack 2 of non-malignant breast tissue, and volume stacks 1, 2, and 3 of malignant breast tissue. The signal intensity values were normalized to the maximum intensity recorded within each respective channel, expressing all values as a proportion of the channel’s maximum. For each stack, DAPI signal intensity and SHG signal intensity were plotted against tissue depth on a single graph (Figure 4, Figure 5). By comparing the trend of the DAPI signal curve to that of the SHG signal, which is only affected by scattering of the excitation light due to intrinsic tissue properties, and remains unaffected by dye penetration, differences in signal attenuation were used to assess dye penetration quality.

**Figure 1:**
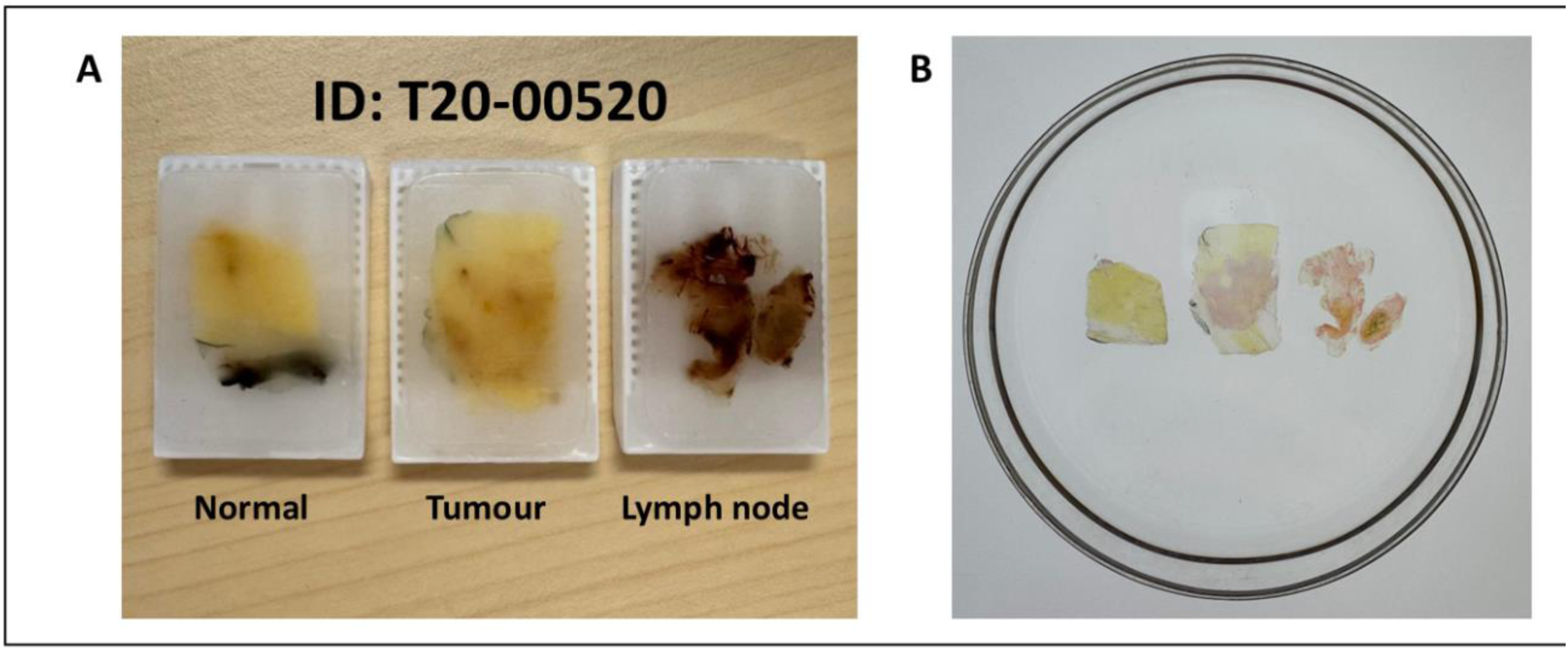
A) FFPE breast and lymph node tissue blocks from patient 1 prior to processing, representing normal breast tissue, tumor (IDC), and axillary lymph node tissue; B) progress picture of cleared and labeled samples

**Figure 2:**
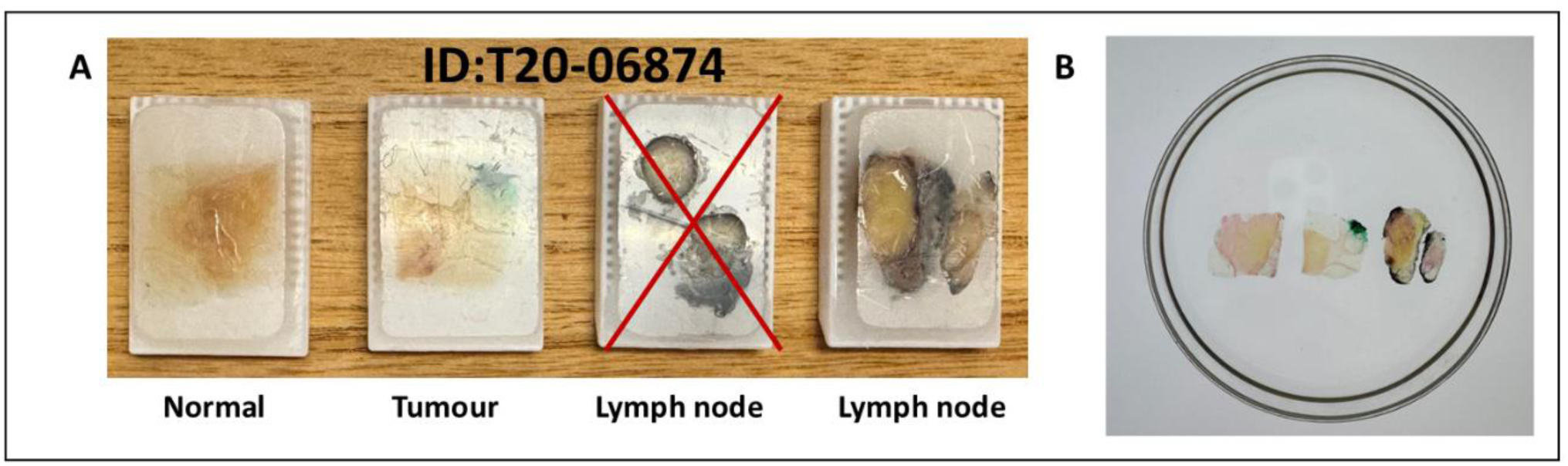
A) FFPE breast and lymph node tissue blocks from patient 2 prior to processing, representing normal breast tissue, tumor (IDC), and axillary lymph node tissue; B) progress picture of cleared and labeled samples

**Figure 3:**
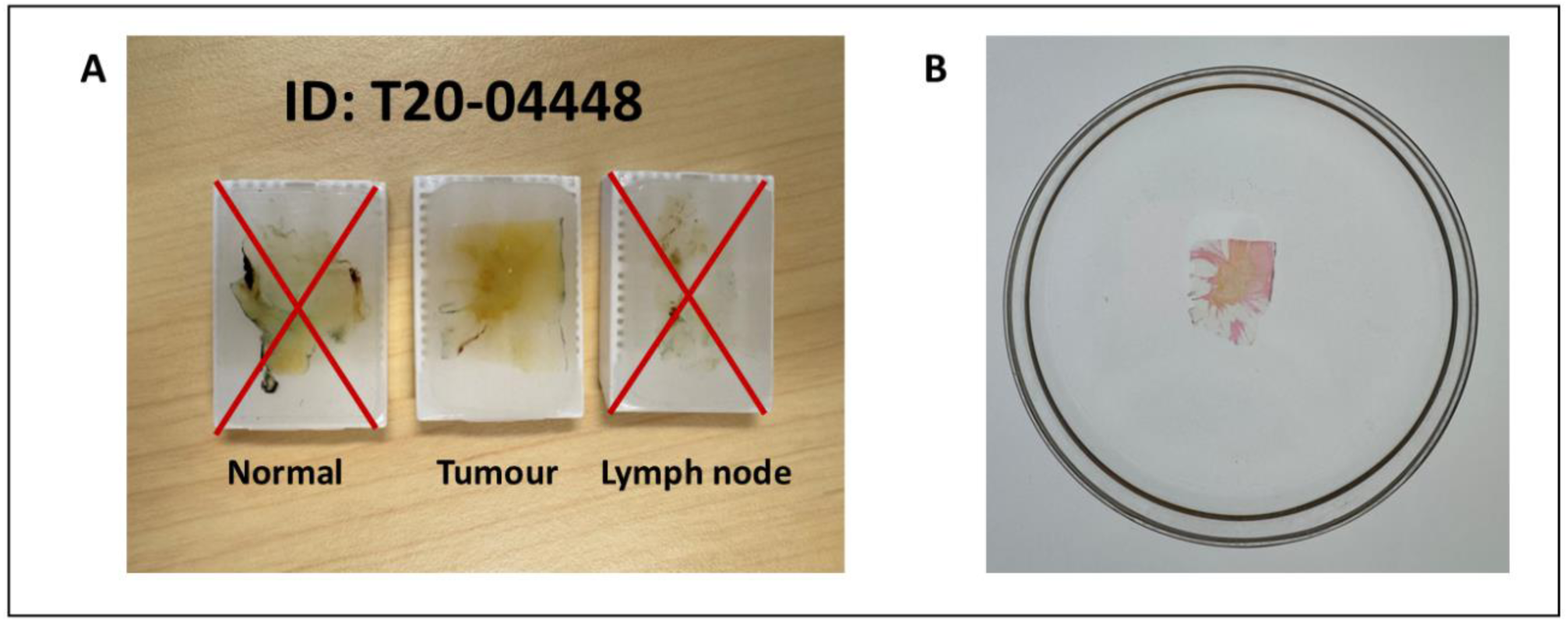
A) FFPE malignant breast tissue block from patient 3 prior to processing, representing normal breast tissue, tumor (IDC), and axillary lymph node tissue; B) progress picture of cleared and labeled samples

**Figure 4:**
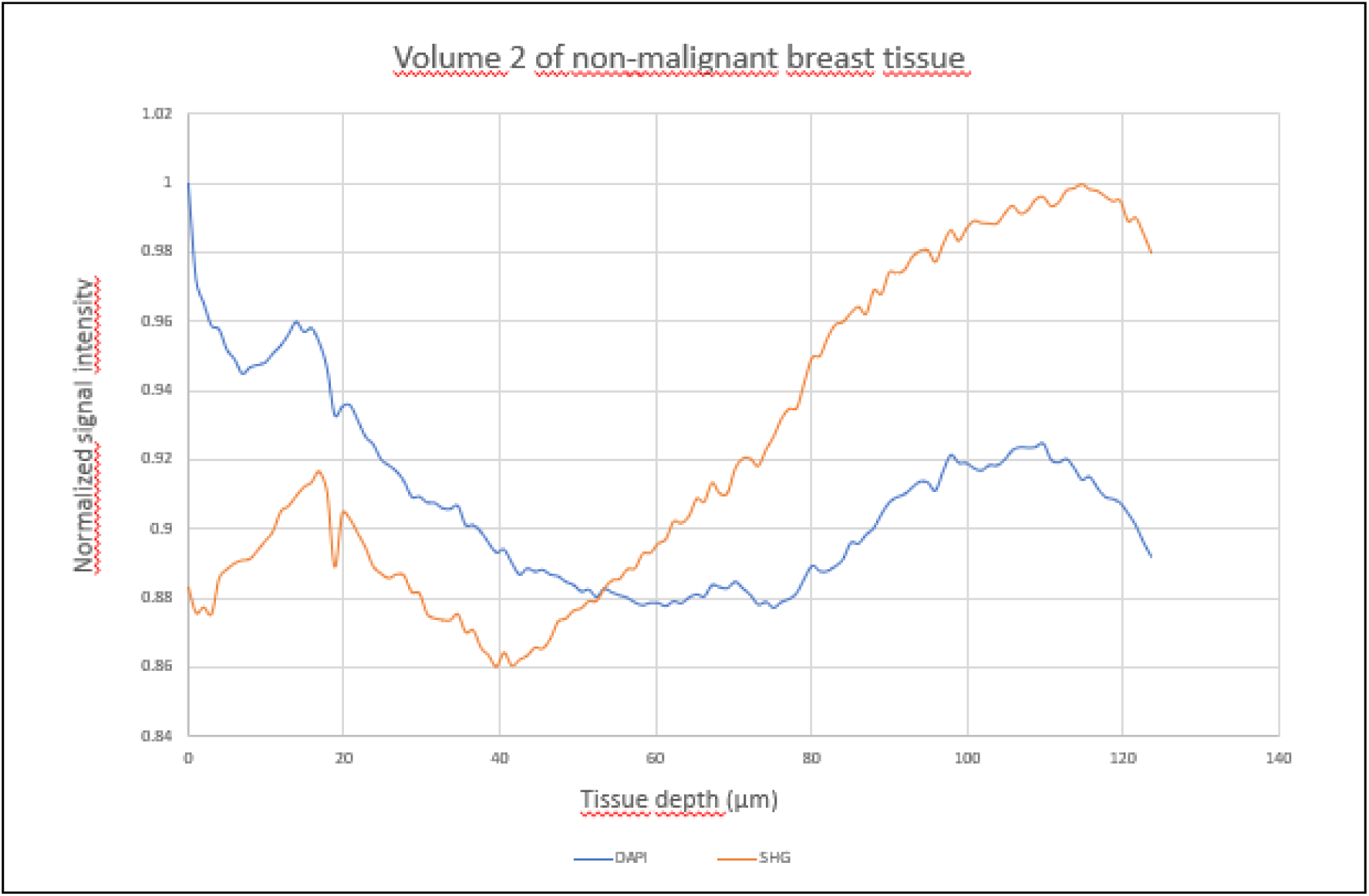
Normalized signal intensity across tissue depth in cleared non-malignant breast tissue (volume 2**)\*** *Depth profile showing normalized signal intensity for nuclear staining (DAPI, blue) and collagen (SHG, orange) across the z-axis of volume 2 from non-malignant breast tissue. Signal values were normalized to each channel’s maximum intensity to allow comparison of depth-dependent signal behavior.

**Figure 5:**
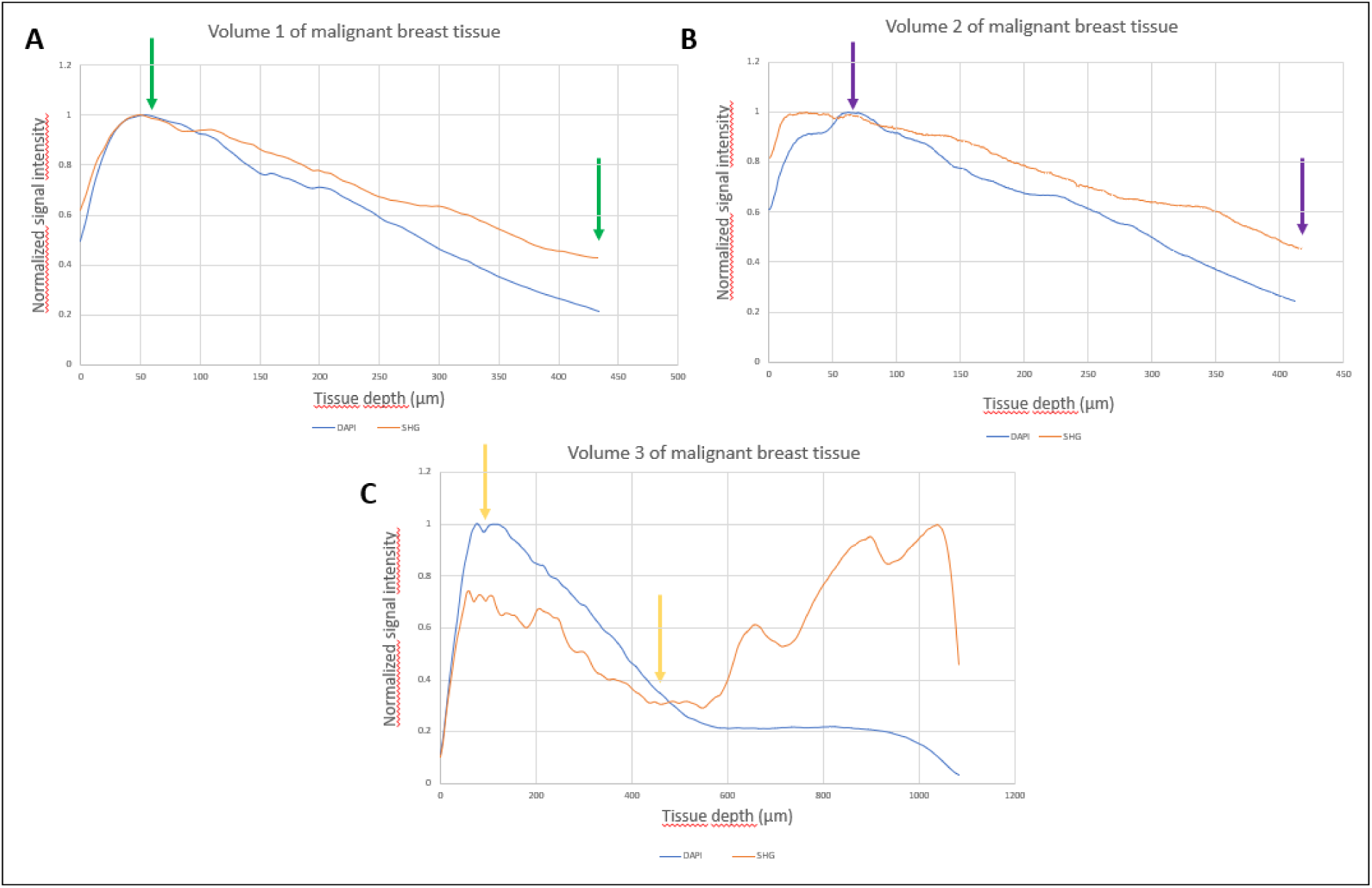
Normalized signal intensity across tissue depth in cleared malignant breast tissue (volume 1(a), volume 2 (b) and volume 3 (c))*

#### 2.5.3 Quantitative Analysis of Tissue Architecture

Quantitative image analysis was performed on the selected frames (Table 2) from previously mentioned volume stacks and images acquired from sample 1 (grade I IDC) malignant breast tissue, adjacent non-malignant breast tissue, and matched lymph node tissue (Figures 6-9).

**Figure 6:**
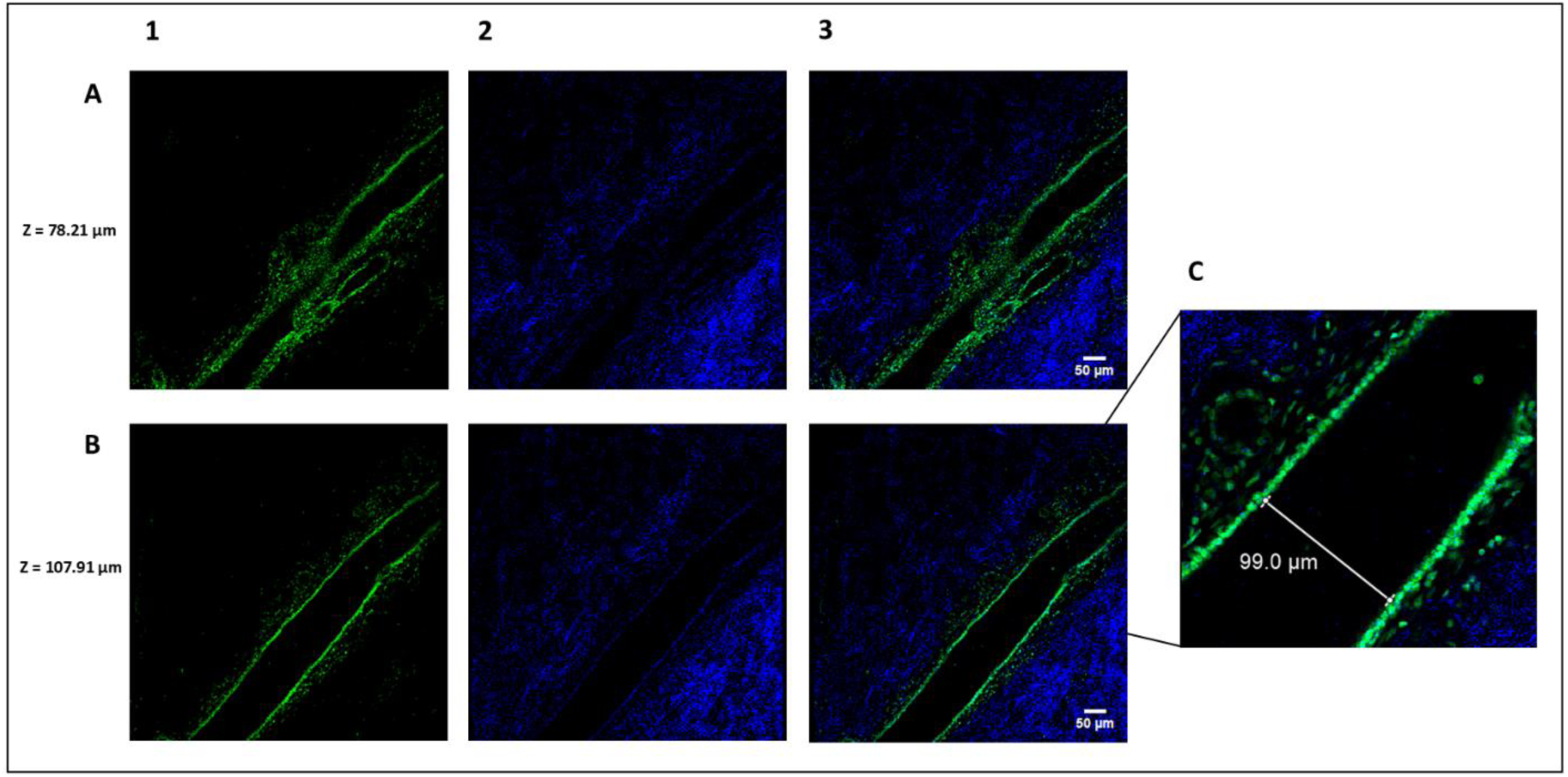
Vascular structure in cleared non-malignant breast tissue visualized by two-photon microscopy; A, B) z-plane images from volume 1 showing a tubular DAPI+ structure (green) with a central lumen, surrounded by SHG+ collagen matrix (blue), at depths of 78.21 µm (A) and 107.91 µm (B); C) a zoomed in view highlighting the hollow lumen, measuring approximately 99.0 µm in diameter * *DAPI = nuclear stain, SHG+= second harmonic generation (collagen); Columns: (1) DAPI nuclear channel, (2) SHG collagen channel, (3) Merged image; Scale bar: 50 µm, consistent across all panels in A and B.

**Figure 7:**
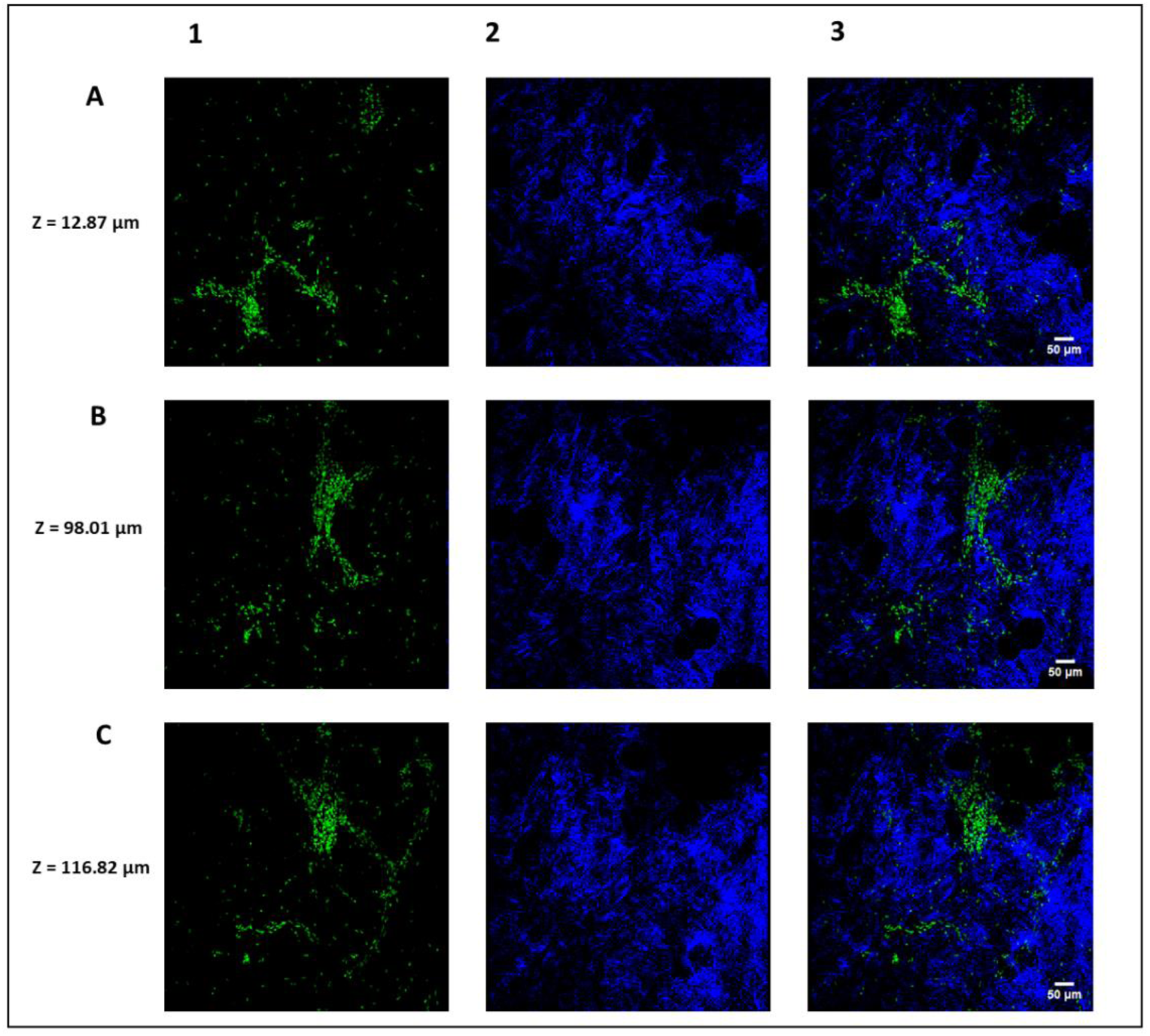
DAPI+ cell clusters and stromal architecture in volume 2 of cleared non-malignant breast tissue, (A-C) Representative z-plane images at depths of 12.87 µm, 98.01 µm, and 116.82 µm, showing dense DAPI+ (green) cell clusters embedded in a structured SHG+ (blue) collagen matrix. Dark signal-void areas are visible throughout the field of view, consistent with adipocyte morphology. The DAPI+ clusters exhibit partial radial or branched arrangements and are frequently adjacent to dense stromal fibers* *DAPI = nuclear stain, SHG+= second harmonic generation (collagen); Columns: (1) DAPI nuclear channel, (2) SHG collagen channel, (3) Merged image; Scale bar: 50 µm, consistent across all panels in A and B.

**Figure 8:**
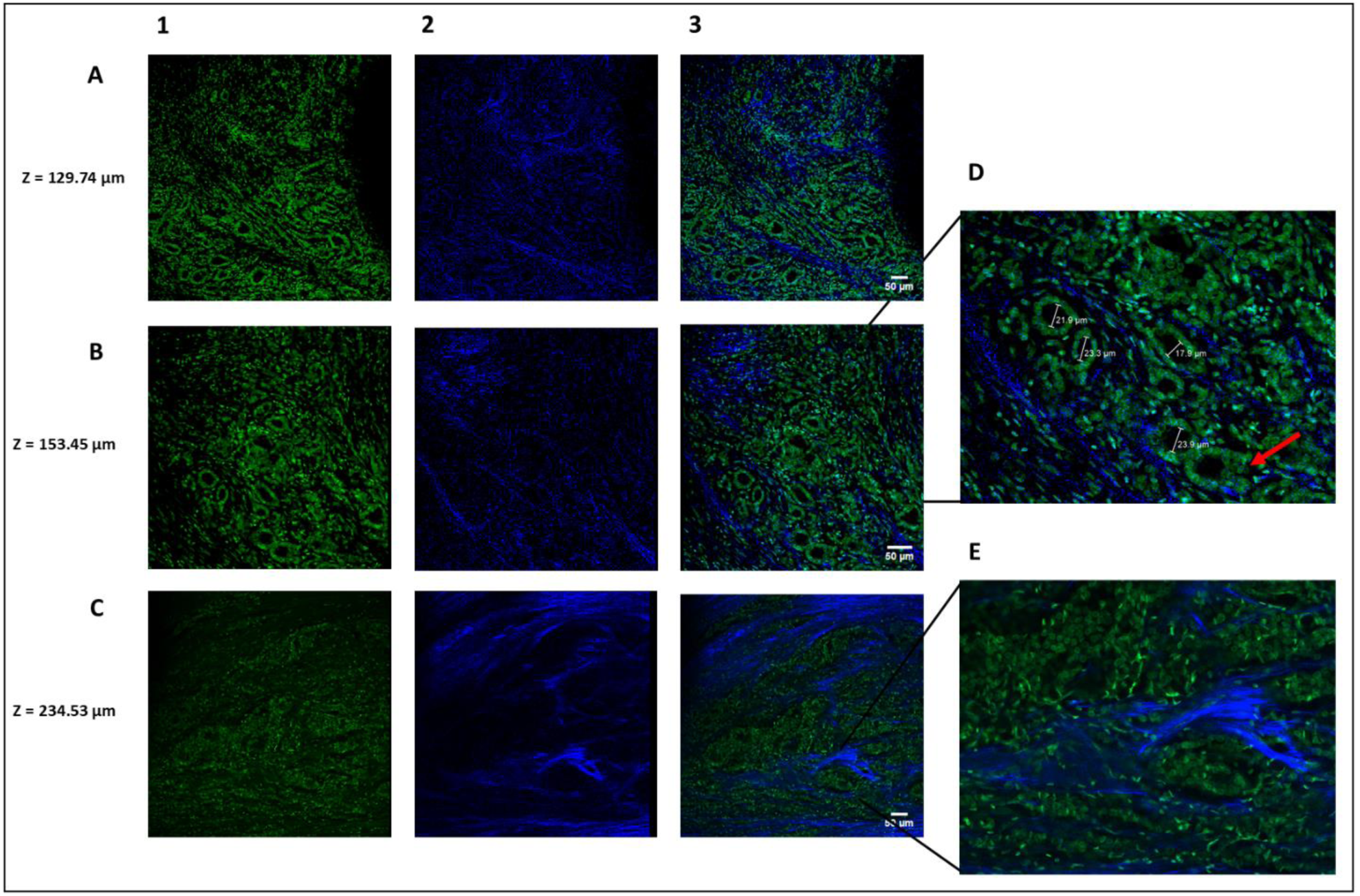
Structural features of malignant breast tissue across three volumes reveal preserved and altered TDLU morphology; A-C) Representative z-planes from volumes 1, 2, and 3 of cleared malignant breast tissue showing DAPI+ epithelial structures (green) and SHG+ collagen matrix (blue); D) Zoomed in view of volume 2 showing DAPI+ TDLU profiles with duct diameters of roughly 15–25 µm; red arrow indicates a disrupted glandular structure; E) Zoomed in view of volume 3 showing increased nuclear pleomorphism and collagen remodeling, with denser and more disorganized SHG+ fibers* *DAPI = nuclear stain, SHG+= second harmonic generation (collagen); Columns: (1) DAPI nuclear channel, (2) SHG collagen channel, (3) Merged image; Scale bar: 50 µm, consistent across all panels in A and B.

**Figure 9:**
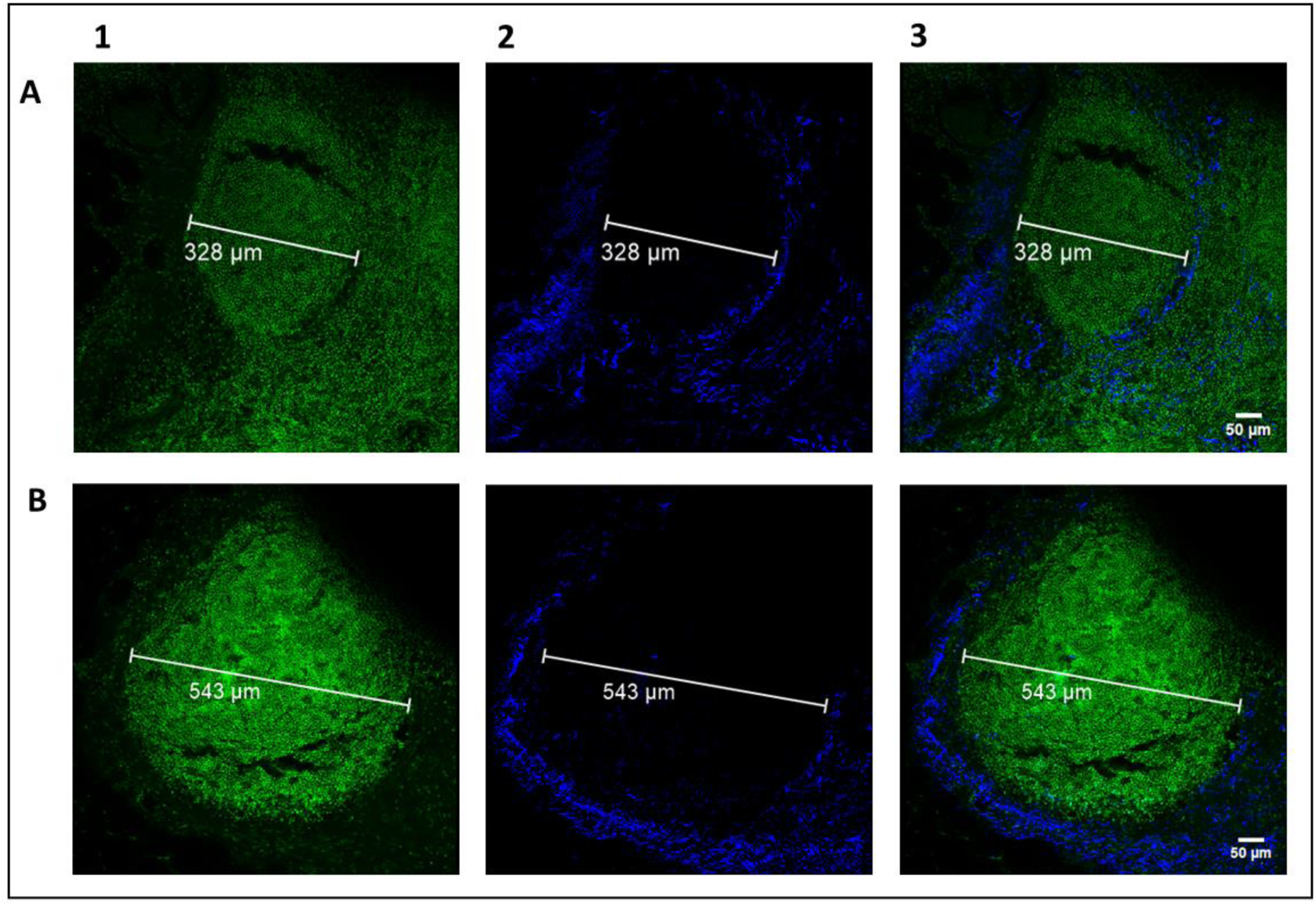
Structural imaging of intact lymphoid follicles in cleared axillary lymph node tissue; A) Z-projected two-photon image across 144.71 µm tissue depth showing a circular lymphoid follicle (328 µm in diameter), B) Single-plane image from the same tissue showing a larger follicle (543 µm in diameter). DAPI+ cells (green) are densely packed and uniform in size and shape, while the SHG+ signal (blue) forms a peripheral collagen rim* *DAPI = nuclear stain, SHG+= second harmonic generation (collagen); Columns: (1) DAPI nuclear channel, (2) SHG collagen channel, (3) Merged image; Scale bar: 50 µm, consistent across all panels in A and B.

**Table 2:**
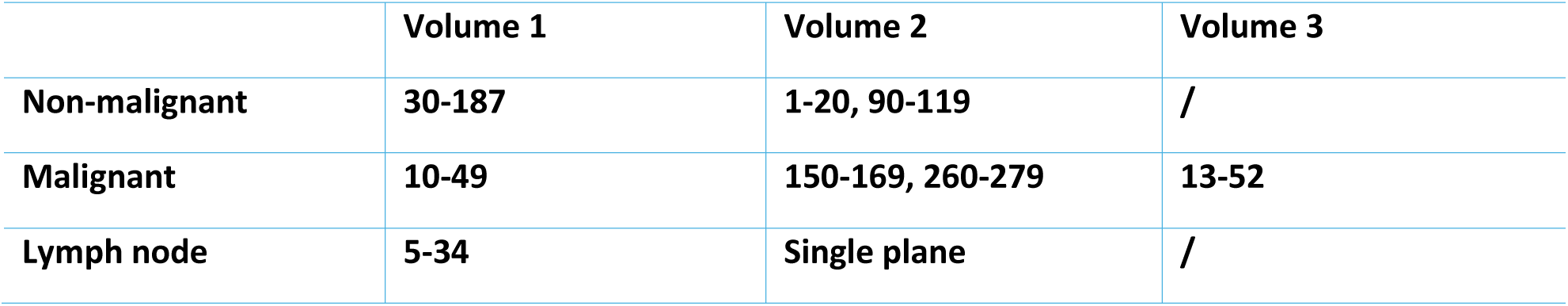
Imaged frames from each of the analyzed volumes selected for quantitative analysis of tissue architecture.

To estimate the mean cell size, total cellular area, and cell count, grayscale images from the nuclear staining channel were processed in FIJI. Images were first converted to 8-bit and subjected to background subtraction using the rolling ball algorithm (radius = 200 pixels) to reduce uneven illumination. Cell regions were segmented using the RenyiEntropy thresholding method, producing a binary image. The segmented cell area and percentage area coverage were then extracted by measuring the area limited to the threshold. The number of individual cells was estimated using the “Find Maxima” function with a prominence threshold set above 10. In this context, prominence refers to the minimum intensity difference between a local maximum (cell nucleus) and its surrounding pixels, helping to distinguish true peaks (i.e., cells) from background noise. The mean cell area was calculated by dividing the total cellular area by the estimated number of cells.

To assess collagen content, the SHG channel, corresponding to extracellular matrix components, primarily collagen, was converted to 8-bit grayscale and background-subtracted (rolling ball radius = 200). Collagen rich regions were segmented using RenyiEntropy thresholding, and the resulting binary mask was used to measure the total collagen area and the area fraction relative to the field of view.

Additionally, the diameters of specific structures such as ducts, lobules, vessels, and lymphoid follicles were manually measured using LAS AF software.

#### 2.5.4 Collagen organization analysis

Collagen fiber organization was analyzed for the SHG channel of the three selected volume stacks acquired from patient 1 (grade I IDC) malignant and adjacent non-malignant breast tissue. Orientation analysis was performed using the OrientationJ plugin and its Distribution function, with a local window (σ) of 2 pixels and Cubic Spline gradient, in FIJI. The acquired data were exported into Excel where the mean number of pixels per slide corresponding to each fiber alignment angle (-90° to +90°) was calculated and used to visualize angular distribution and identify dominant orientation peaks (Figures 10b,11b,12b).

**Figure 10:**
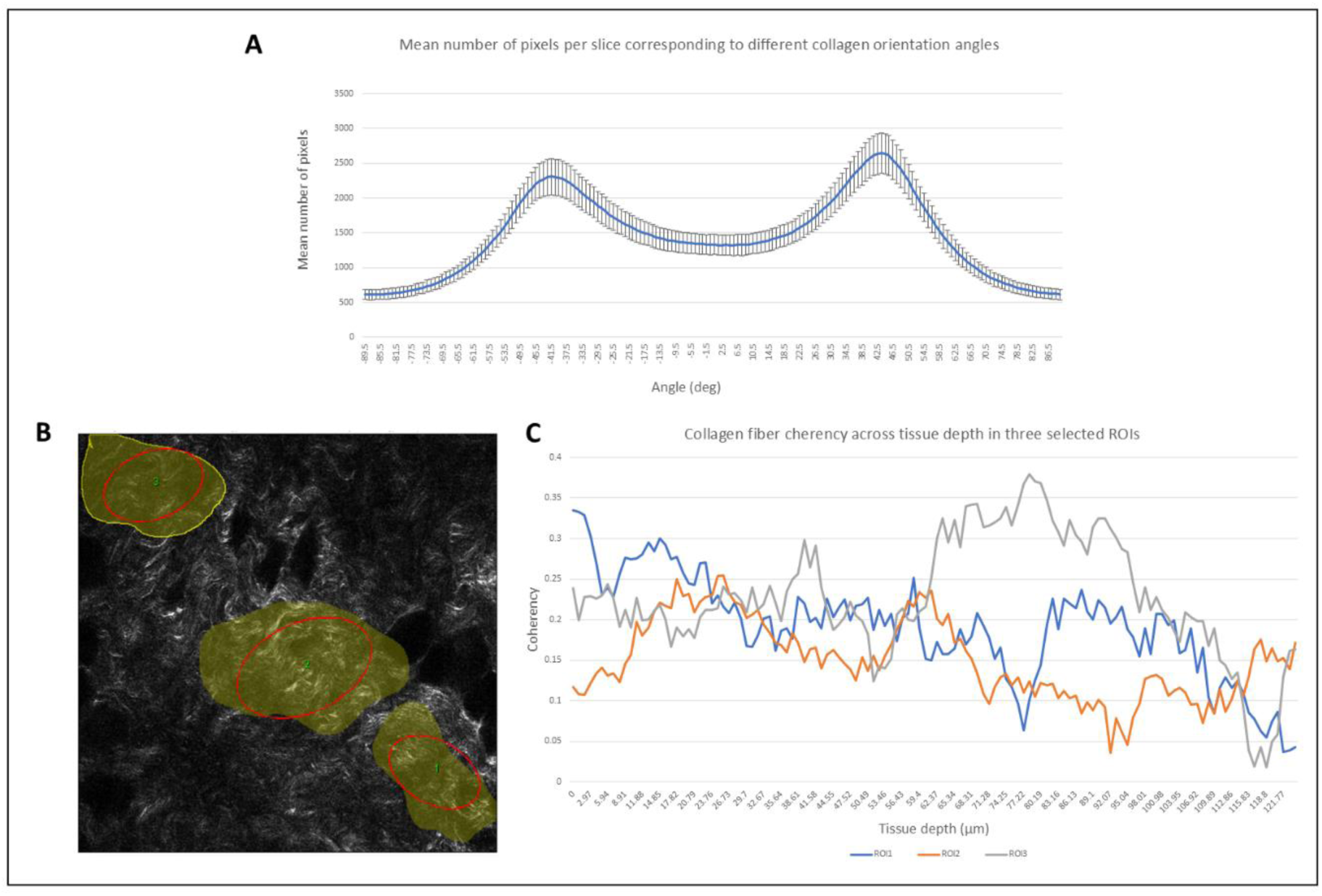
Collagen fiber orientation and coherency in volume 1 of cleared non-malignant breast tissue; A) Histogram showing a bimodal distribution of collagen fiber orientation angles across all slices, with peaks centered around ±45°, indicating a consistent angular organization; B) Representative SHG image of collagen fibers with selected ROIs (yellow) used for coherency analysis; C) Coherency values plotted across tissue depth for three selected ROIs, showing overall low-to-moderate alignment, consistent with a disordered collagen matrix* *SHG = second harmonic generation (collagen signal); ROI = region of interest.

**Figure 11:**
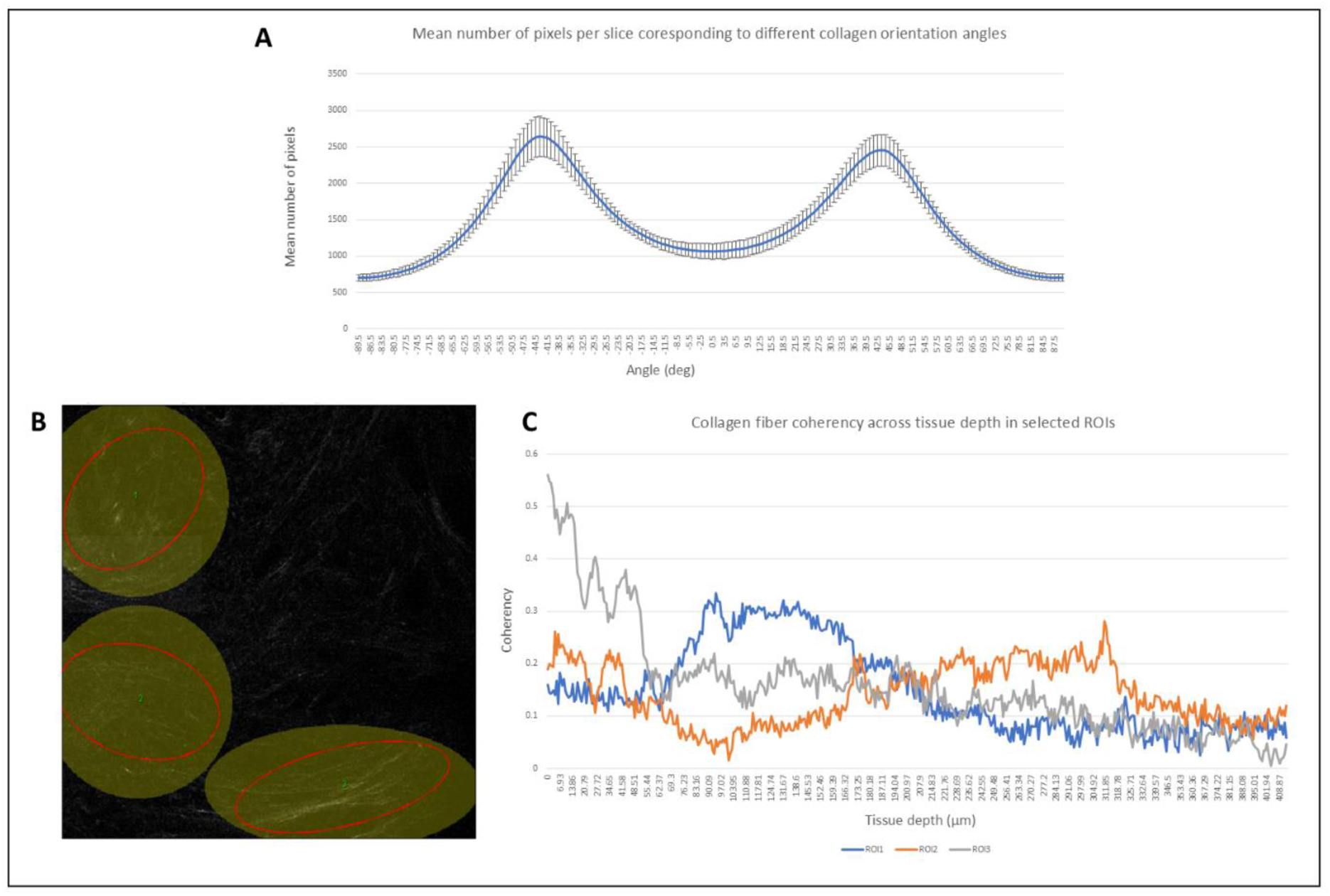
Collagen fiber orientation and coherency in volume 2 of cleared malignant breast tissue; A) Histogram showing a bimodal distribution of collagen fiber orientation angles across all slices, with peaks centered around ±45°, indicating a consistent angular organization; B) Representative SHG image of collagen fibers with selected ROIs (yellow) used for coherency analysis; C) Coherency values plotted across tissue depth for three selected ROIs, showing overall low-to-moderate alignment, consistent with a disordered collagen matrix* *SHG = second harmonic generation (collagen signal); ROI = region of interest.

To quantify collagen alignment, multiple ROIs were manually selected (Figures 10c, 11c, 12c) and evaluated for each of the three tissue volume stacks using OrientationJ > Measure, with σ = 2. The resulting coherency values, which range from 0 (completely random orientation) to 1 (perfect alignment), were extracted for each ROI and plotted as a function of tissue depth in Excel (Figures 10D, 11D, 12D).

## Results

### 3.2 Deparaffinization and Tissue Clearing

Tissue processing was successful for malignant, non-malignant, and lymph node samples from patients 1 and 2, as well as for the malignant tissue of patient 3. However, both the non-malignant and lymph node samples from patient 3 sustained extensive structural damage during deparaffinization, making them unsuitable for further clearing and analysis. These samples were therefore excluded from the study.

Among the successfully processed samples, tissue integrity was largely preserved. A minor detachment of the bottom portion of the non-malignant breast tissue from patient 1 was observed, but this did not compromise the remainder of the sample. Nearly complete optical transparency was achieved across all included specimens (Figures 1B, 2B, 3B). Residual surgical dye was observed at the periphery of the lymph node tissue from patient 2 (Figure 2B), but this did not interfere with subsequent imaging.

### 3.2 Tissue Shrinkage

The mean surface area before tissue processing was 317.802 ±135.612 mm², which decreased to 296.510 ±136.186 mm² post-treatment. This represents a mean reduction of 21.292 ±13.878 mm², accounting for roughly 6.7% of the original surface area. The null hypothesis, stating that there is no difference in surface area before and after processing (mean pre = mean post), was tested against the alternative hypothesis that tissue surface area decreases significantly (mean pre > mean post). A one-tailed paired samples t-test yielded a p-value of < 0.001, demonstrating statistically significant tissue shrinkage during processing. However, in practice, the tissue shrinkage percentage is complies with that in other successful, similar protocols [19]. Additionally, prior to the processing, tissue thickness ranged from approximately 0.9 to 1.6 mm and decreased slightly after treatment to 0.8 to 1.4 mm.

### 3.3 DAPI Dye Penetration

#### 3.3.1 Non-malignant breast tissue

In volume stack 2 of non-malignant breast tissue (Figure 4), the normalized signal intensity profiles of the DAPI and SHG channels display a similar pattern of increase and decrease over the same depth intervals. A notable exception occurs between approximately 40 and 55 µm, where the SHG signal begins to rise after reaching its minimum, while the DAPI signal intensity continues to decline, resulting in a brief divergence. Shortly beyond the point of intersection, both curves start exhibiting an upward trend, with the SHG signal increasing at a slightly higher rate of increase than DAPI. The overall alignment of the two curves across most of the imaged depth suggests that the nuclear dye signal curve closely follows that of the tissue collagen structure, as reflected by SHG. This correspondence supports the conclusion that dye penetration was effective throughout the most of the imaged volume of 123.5 µm.

#### 3.3.2 Malignant breast tissue

In all three imaged volumes of malignant breast tissue, the normalized signal intensity profiles of the DAPI and SHG channels demonstrate similar increase/decrease trends. In each case, both curves reach their peak intensity at approximately 60 µm of tissue depth, followed by a near-linear decline, as indicated with green (Figure 5A), purple (Figure 5B), and yellow (Figure 5C) arrows. In volume stacks 1 and 2, which extend to 434.08 µm and 411.99 µm, respectively, the similar rate of decrease in both DAPI and SHG signals suggests effective dye penetration throughout the imaged depth. In contrast, volume 3 extends to a depth of 1082.7 µm. While the DAPI signal plateaus at a low intensity between 600 and 1000 µm, the SHG signal diverges markedly, exhibiting a sharp increase and reaching its maximum at 1037.80 µm. This divergence indicates that, while structural features remain detectable at greater depths, nuclear dye penetration was limited, failing to label deeper regions as effectively as in the shallower stacks.

### 3.4 Quantitative Analysis of Tissue Architecture

#### 3.4.1 Non-malignant breast tissue

2P imaging of volume 1 from the non-malignant breast tissue (Figure 6) revealed an elongated tubular structure, extending across 155.43 µm in depth, with a clearly defined, dark, and hollow central lumen, measuring approximately 99 µm in diameter (Figure 6C). The luminal surface was lined by a thin, continuous single layer of DAPI+ cells (green), with the mean surface area of 16.832±1.537µm², consistent in appearance with flattened squamous endothelial cells forming the capillary wall [21] (Figures 6A1 and 6B1) The structure was surrounded by a dense sheet-like, SHG+ area (blue), making up, on average, 18.353±0.825% of the imaged frame, and was consistent with the collagen-rich extracellular environment of interlobular stroma [22] (Figures 6A2a and 6B2). The structural organization and stromal embedding suggest intact vascular architecture and preserved physiological features, supporting the effectiveness of the clearing and staining protocol in maintaining native tissue morphology.

Figure 7, obtained by imaging volume 2 of the non-malignant breast tissue, showed several dense clusters of DAPI+ cells (green) embedded within a highly structured SHG+ (blue) collagenous matrix (on average 40.545±1.458% of the imaged area). The average area of individual DAPI+ cells across the entire imaged volume was 61.547 ± 4.391 µm². Scattered throughout the field of view were large, dark, signal-void regions with diameters ranging from approximately 50 to 100 µm. These features likely correspond to adipocytes, the most abundant features in breast tissue [22]. The spatial arrangement of the DAPI+ clusters showed partial radial or branched organization and frequent proximity to dense stromal fibers. This morphology may suggest structural similarity to TDLUs, however, several features required for definitive identification of TDLUs were not observed. Specifically, no distinct acinar formations, terminal ductal lumina, or the characteristic bilayered epithelial pattern [23], were detected. One plausible alternative explanation, based on the size of the cells and their slightly irregular shape, is that these DAPI+ clusters represent immune cell aggregates, such as lymphocytes [24].

#### 3.4.2 Malignant breast tissue

In volumes 1 (Figure 8A-1,2,3) and 2 (Figure 8B-1,2,3) of malignant breast tissue, transverse sections of DAPI+ TDLU structures (green) were observed (Figures 8A1 and 8B1), with most ductal or acinar profiles measuring approximately 15 to 25 µm in diameter (Figure 8D). These glandular structures appeared largely intact, with only a few displaying signs of structural disruption (Figure 8D, red arrow). The epithelial cells lining the ducts exhibited relatively uniform morphology, with minimal variation in nuclear size and shape, consistent with a low degree of nuclear pleomorphism. The surrounding SHG+ connective tissue (blue) was loosely organized, occupying an average of 26.107 ± 0.835% and 12.647 ± 3.154% of the imaged area in volumes 1 and 2, respectively (Figures 8A2 and 8B2). In contrast, volume 3 (Figure 8C) displayed a more altered tissue architecture. Tubular structures were poorly defined, and the tissue exhibited clear architectural distortion (Figure 8C1). Nuclear morphology was more heterogeneous, with more pronounced variation in nuclear size and shape (Figure 8E), indicative of increased nuclear pleomorphism. The extracellular matrix was denser, with thick, infiltrating collagen fibers making up 34.191 ± 2.579% of the imaged area (Figure 8C2), consistent with a desmoplastic stromal response [25].

Across all three volumes, the mean nuclear area was 24.737 µm², a value within the range typically observed in healthy breast tissue [11]. However, the relatively large standard deviation (8.891 µm²) likely reflects local variability in nuclear size, particularly in regions with pleomorphic features. Collectively, the morphology observed across these volumes is consistent with grade I invasive ductal carcinoma (IDC) [5], while also highlighting the intra-tumoral heterogeneity in tissue structure and cellular morphology.

#### 3.4.3 Lymph node tissue

The imaging of 144.71 µm of tissue depth within the axillary lymph node tissue (Figure 9A) and a single plane image acquired from the same tissue (Figure 9B) showed large, circular structures, 328 to 543 μm in diameter. The green, nuclear DAPI channel revealed densely packed cells throughout the structure (Figure 9A1 and 9B1). These cells were uniformly small, averaging at 9.659±4319 µm² surface area, and had a consistently rounded shape. The blue SHG signal (collagen) outlined the structure, forming a thin, peripheral rim around the entire cell-dense area (on average 6.117±2.099% of the imaged area), and was absent in the interior (Figures 9A2 and 9B2). The architecture, nuclear morphology, and stromal organization observed in the two-photon image are consistent with that of a lymphoid follicle [26] a key structure in the cortex of lymph nodes. Additionally, no signs of follicular disruption, loose epithelium, pleomorphic nuclei, or gland-like structures confirm this is non-metastatic, structurally intact lymphoid tissue [27].

### 3.5 Collagen organization analysis

Collagen fiber orientation analysis revealed a distribution similar to bimodal distribution centered around ±45° in both volume 1 of non-malignant breast tissue (Figure 10A) and volume 2 of malignant breast tissue (Figure 11A), suggesting consistent angular organization across tissue depth. In volume 3 of malignant tissue (Figure 12A), the distribution was slightly shifted toward a dominant 45° orientation, potentially reflecting the early stages of collagen remodeling.

**Figure 12:**
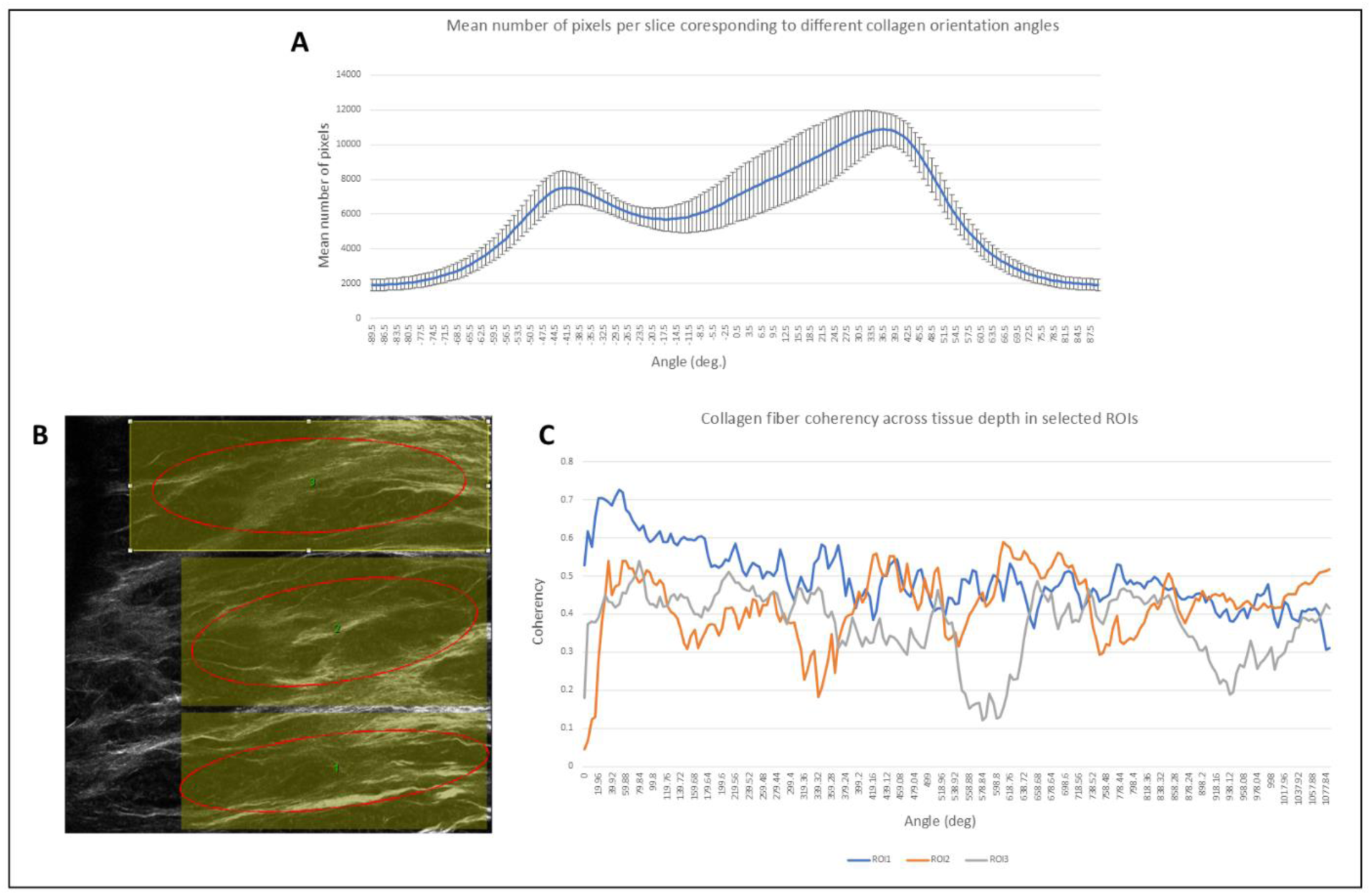
Collagen fiber orientation and coherency in volume 3 of cleared malignant breast tissue; A) Histogram showing a shifted bimodal distribution of collagen orientation angles, with one dominant peak near 45°, suggesting directional remodeling, B) Representative SHG image of collagen fibers with selected ROIs (yellow) used for coherency analysis; C) Coherency values plotted across tissue depth for three selected ROIs, showing overall moderate alignment, consistent with the remodeling of collagen matrix* *SHG = second harmonic generation (collagen signal); ROI = region of interest.

Fiber coherency values in volume 1 of non-malignant (Figure 10C) and volume 2 of malignant (Figure 11C) tissue were relatively low and variable, with most values ranging between 0.1 and 0.3, indicative of disordered collagen alignment. In contrast, volume 3 of malignant tissue (Figure 12C) exhibited somewhat higher coherency values, predominantly between 0.3 and 0.6, suggesting a more organized and aligned collagen architecture.

Together, these results indicate that both non-malignant and low-grade malignant breast tissue (grade I IDC) display low to moderate collagen fiber alignment, despite consistent orientation angles. In regions exhibiting more prominent cancer morphology, seen in volume 3 of malignant tissue (Figure 8C), there is evidence of increased collagen remodeling, manifested as higher fiber alignment and more uniform orientation patterns.

## Discussion

The present study demonstrates the feasibility and advantages of combining the MASH clearing protocol with 2P imaging for large FFPE human breast and lymph node tissue samples. To our knowledge, this represents the first successful adaptation of MASH for FFPE samples of this scale. The protocol was modified to incorporate ECi as a RIMS and dye combinations were adapted. High tissue transparency and good structural preservation were achieved, even in millimeter-thick samples.

Tissue shrinkage analysis, based on 2D surface area measurements before and after processing, revealed an average reduction of 6.7%. While statistically significant (p < 0.001), this reduction is consistent with shrinkage values reported in other clearing protocols [19]. Due to non-uniform sample thickness and imprecise depth estimation, z-axis shrinkage could not be reliably quantified, representing a limitation. Future studies using digital 3D volume tracking of each processing stage may enable more accurate assessment of volume shrinkage.

Nuclear dye penetration was generally successful, with DAPI signal intensity curves closely following the SHG signal intensity curve behaviors up to a 600 µm tissue depth (Figure 4, Figure 5). Beyond this point, DAPI signal decline was observed (Figure 5C), potentially due to reduced staining efficacy. However, these results must be interpreted cautiously, as alternative factors, such as localized cell death, could also contribute to reduced signal. Validation on a larger sample set with varied tissue thicknesses will be necessary to disentangle these effects and optimize labeling protocols accordingly.

The presented protocol preserved cytoarchitecture and collagen integrity, supporting compatibility with dual-channel multiphoton imaging. These findings position the method as a promising foundation for volumetric pathology workflows in research contexts.

Traditional breast cancer diagnostics rely on 2D histological sections limited to 5 µm thickness, inherently restricting spatial interpretation. In contrast, 2P imaging of optically cleared samples enabled visualization of cellular and stromal features over depths up to 0.6 mm and 1 mm, respectively (Figure 5C). This allowed detailed exploration of structures such as TDLUs (Figure 8A1, Figure 8B1), adipocytes (Figure 7B1-3), blood vessels (Figure 6), and lymphoid follicles in their intact 3D spatial context (Figure 9). Importantly, the increased depth facilitated detection of local architectural disruption and nuclear pleomorphism across tissue layers (Figure 8C1-3), which can more easily be missed in conventional single-slice analysis. This is particularly relevant in light of increasing interest for intratumoral heterogeneity research.

Analysis of collagen organization further highlights the strengths of volumetric imaging. Despite both malignant volumes being derived from the same grade I IDC sample, morphologically distinct regions revealed notable differences in collagen features. Structurally preserved areas, whether non-malignant or less disrupted malignant tissue, exhibited loosely organized SHG+ fibers with low coherency values (Figures 7A2, 7B2, 7C2, 8A2, 8B2, 10C, and 11C). In contrast, morphologically disrupted regions showed increased fiber density (Figure 8C2), alignment (Figure12C), and shifts in orientation distribution (Figure 12A), suggesting that the collagen remodeling of the tissue had begun. These findings underscore the ability of 3D imaging to uncover spatially confined microenvironmental changes, supporting its potential in future tumor-stroma interaction studies.

Beyond tumor analysis, the protocol also proved effective for imaging of matched lymph nodes (Figure 9). Lymphoid follicles appeared structurally intact, with preserved stromal boundaries (Figure 9A2, Figure 9B2) and cellular density (Figure 9A1, Figure 9B1), features indicative of non-metastatic morphology [27]. Additionally, identification of a long DAPI+ endothelial structure with an open lumen in non-malignant breast tissue (Figure 6) further illustrates the preservation of vascular features. These results demonstrate the pipeline’s broader applicability for studying vascular remodeling and immune infiltration in 3D.

Nonetheless, several limitations must be acknowledged. Dye signal attenuation beyond 600 µm in malignant tissue highlights the need for extended or modified labeling protocols, particularly in thicker sections. The small sample size and limited number of imaged volumes restrict generalizability and preclude stratified comparisons. Future work should expand the cohort and include samples across different IDC grades to enable statistically robust comparisons. Additional analyses using other dyes, such as NRI, MG, acridine orange (AO), and fluorescein green (FG), are also warranted. Furthermore, integrating this approach with light-sheet microscopy (LSM) could enable large-scale tissue screening, followed by high-resolution 2P imaging of selected ROIs, thereby forming a scalable platform for future volumetric histopathology. The dye combinations applied in the presented staining protocol (Table 1), allows for the reusing of these tissue samples and applying them to alternative imaging modalities, such as LSM.

## Conclusion

This study establishes the feasibility of adapting the MASH clearing and labeling protocol for high-resolution two-photon imaging of FFPE human breast and lymph node tissues, enabling high-resolution two-photon imaging of tumor architecture, including ECM components across millimeter-scale volumes. Despite some signal attenuation in deeper regions, DAPI dye penetration was largely effective, and collagen integrity remained intact throughout the imaging depth. This allowed detailed visualization of ductal and vascular structures, and stromal organization in their native 3D context. Importantly, differences in collagen density and alignment across regions of the same grade I IDC tissue revealed spatially heterogeneous early remodeling events, reinforcing the importance of volumetric approaches in tumor characterization. The presented pipeline was also applicable to adjacent non-malignant tissues and lymph nodes, supporting its versatility across tissue types. Limitations such as depth-dependent dye penetration and small sample size should be addressed in future work. Integration with LSM and expansion across IDC grades may enable novel, high-throughput volumetric pathology workflows for investigating intratumoral heterogeneity and microenvironmental evolution.

## Funding Information

This work was funded by a research grunt of the University Fund Limburg (awarded to A. Schueth), CoBes24.074.

## Conflict of Interest

## Author Information

## 7.2 Supplementary data

### 7.2.1 2P imaging parameters

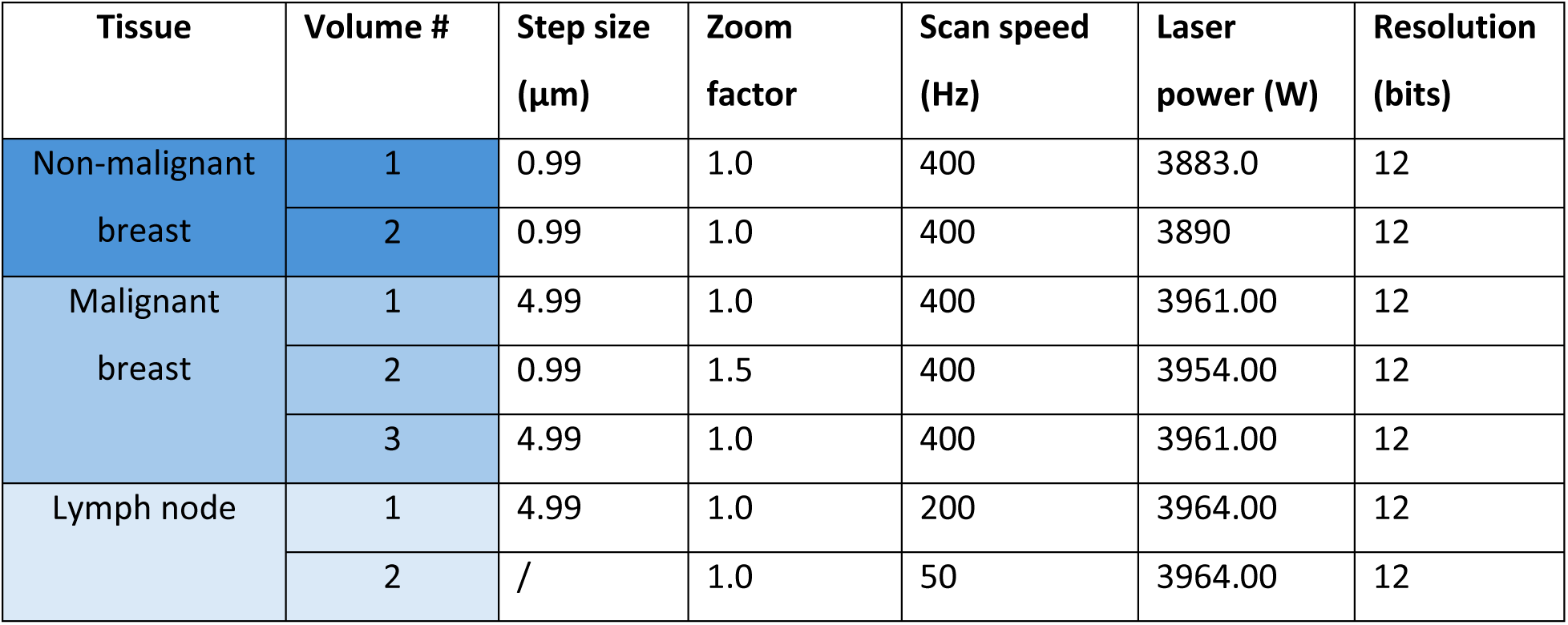

| Tissue | Volume # | Step size (μm) | Zoom factor | Scan speed (Hz) | Laser power (W) | Resolution (bits) |
| --- | --- | --- | --- | --- | --- | --- |
| Non-malignant breast | 1 | 0.99 | 1.0 | 400 | 3883.0 | 12 |
|  | 2 | 0.99 | 1.0 | 400 | 3890 | 12 |
| Malignant breast | 1 | 4.99 | 1.0 | 400 | 3961.00 | 12 |
|  | 2 | 0.99 | 1.5 | 400 | 3954.00 | 12 |
|  | 3 | 4.99 | 1.0 | 400 | 3961.00 | 12 |
| Lymph node | 1 | 4.99 | 1.0 | 200 | 3964.00 | 12 |
|  | 2 | / | 1.0 | 50 | 3964.00 | 12 |

### 7.2.1 Tissue shrinkage analysis in FIJI

**Supplementary Figure 1:**
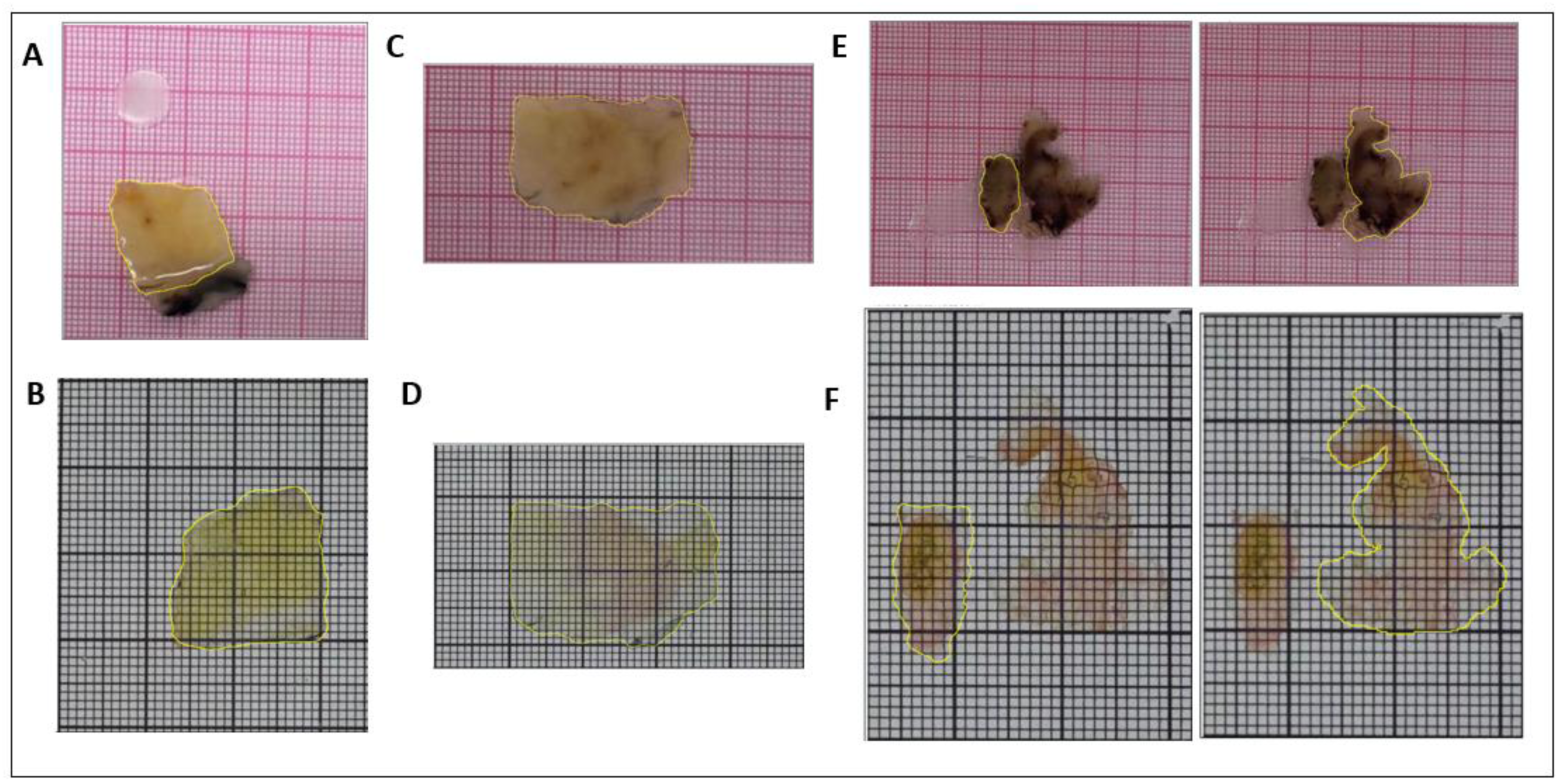
Tissue areas from patient 1 selected for measurements in FIJI; A) non-malignant tissue before processing; B) non-malignant tissue after processing; C) malignant tissue before processing; D) malignant tissue after processing; E) lymph node tissue before processing; F) lymph node tissue after processing

**Supplementary Figure 2:**
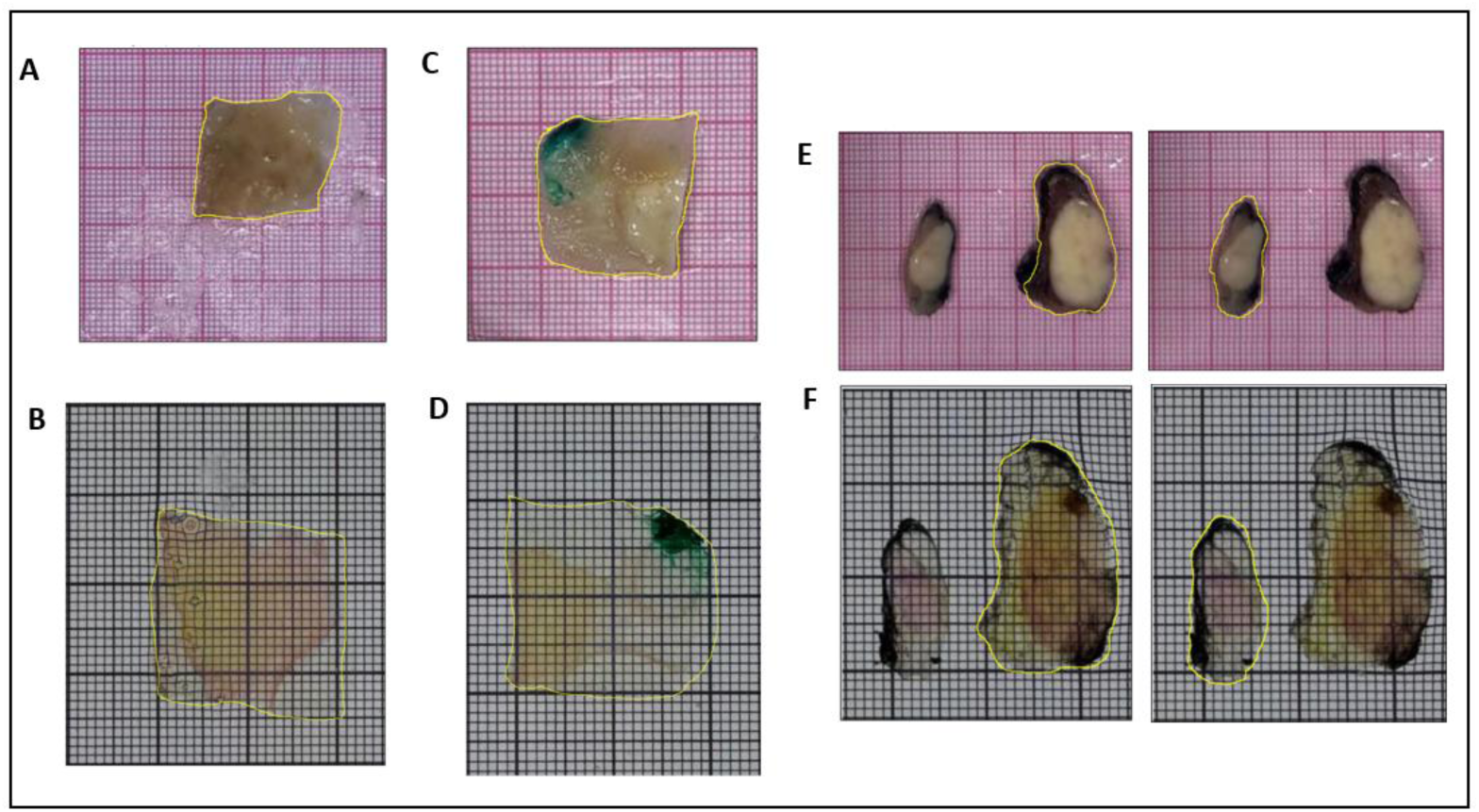
Tissue areas from patient 2 selected for measurements in FIJI; A) non-malignant tissue before processing; B) non-malignant tissue after processing; C) malignant tissue before processing; D) malignant tissue after processing; E) lymph node tissue before processing; F) lymph node tissue after processing

**Supplementary Figure 3:**
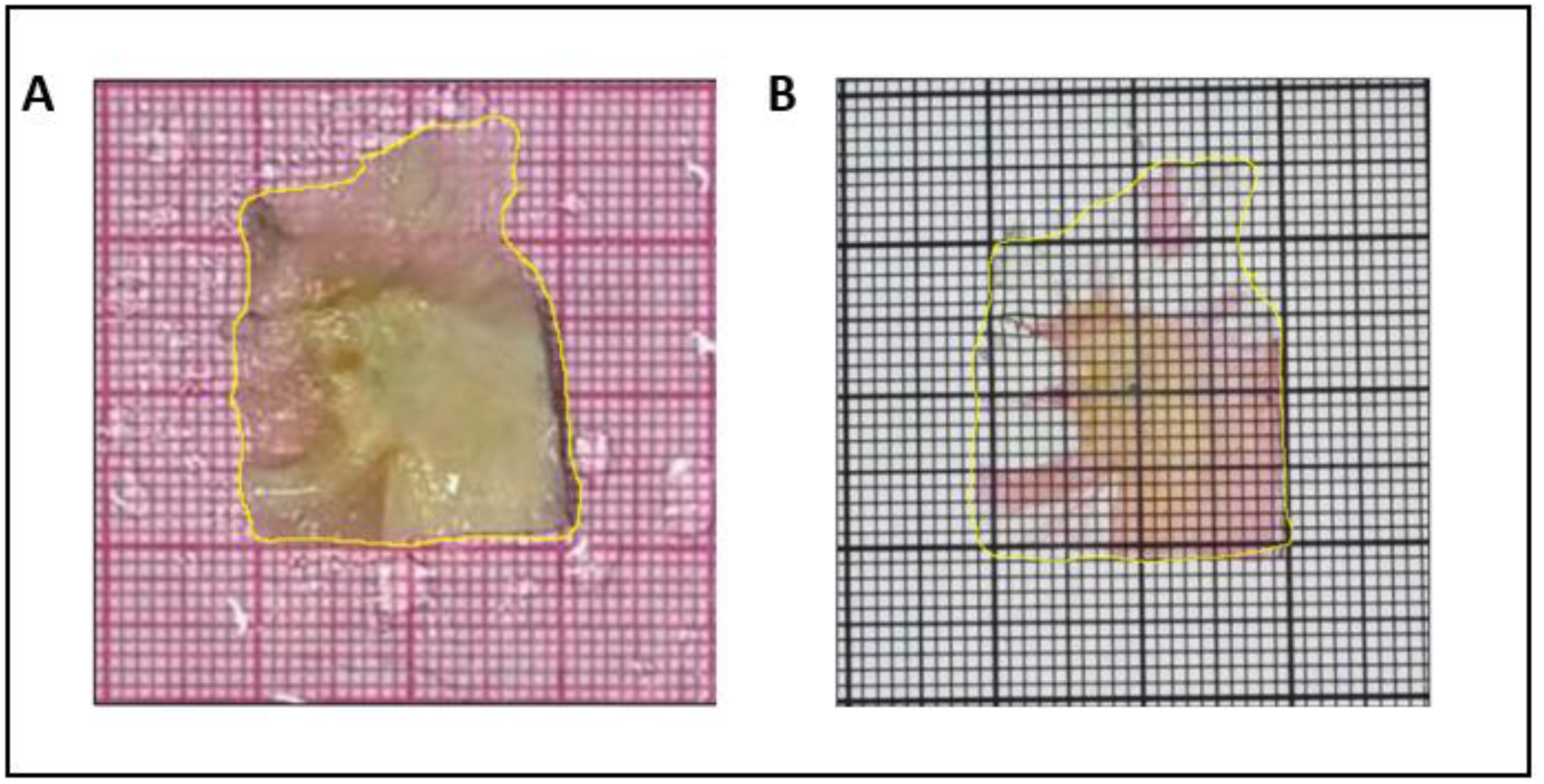
Tissue areas from patient 3 selected for measurements in FIJI; A) malignant tissue before processing; B) malignant tissue after processing

## Notes

### Competing Interest Statement

Declaration of Competing Interest
T. van Nijnatten reports speaker honoraria, institutional grant support and medical advisory board meetings for Bayer and GE Healthcare, not related to the current manuscript. T. van Nijnatten reports medical advisory board for Screenpoint Medical, not related to the current manuscript.

### Summary of Updates

included Albert Bitorina in the author list

